# The erratic and contingent progression of research on territoriality: a case study

**DOI:** 10.1101/107664

**Authors:** Ambika Kamath, Jonathan Losos

**Affiliations:** Department of Organismic and Evolutionary Biology and the Museum of Comparative Zoology, Harvard University, 26 Oxford Street, Cambridge, MA, 02138, USA

**Keywords:** *Anolis*, history, mating system, territorial, polygyny

## Abstract

Our understanding of animal mating systems has changed dramatically with the advent of molecular methods to determine individuals’ reproductive success. But why are older behavioral descriptions and newer genetic descriptions of mating systems often seemingly inconsistent? We argue that a potentially important reason for such inconsistencies is a research trajectory rooted in early studies that were equivocal and overreaching, followed by studies that accepted earlier conclusions at face value and assumed, rather than tested, key ideas about animal mating systems. We illustrate our argument using *Anolis* lizards, whose social behavior has been studied for nearly a century. A dominant view emerging from this behavioral research was that anoles display strict territorial polygyny, where females mate with just the one male in whose territory they reside. However, all genetic evidence suggests that females frequently mate with multiple males. We trace this mismatch to early studies that concluded that anoles are territorial based on limited data. Subsequent research assumed territoriality implicitly or explicitly, resulting in studies that were unlikely to uncover or consider important any evidence of anoles’ departures from strict territorial polygyny. Thus, descriptions of anole behavior were largely led away from predicting a pattern of female multiple mating. We end by considering the broader implications of such erratic trajectories for the study of animal mating systems, and posit that precise definitions, renewed attention to natural history, and explicitly questioning assumptions made while collecting behavioral observations will allow us to move towards a fuller understanding of animal mating systems.

## Introduction

Variation among species in social organization and mating system has long been of interest to naturalists and evolutionary biologists. Why are some species monogamous, others polygynous, and yet others polyandrous? Why do some species exhibit a wide range of reproductive and social behavior? Understanding the selective pressures driving such variation requires quantifying the extent to which different behaviors lead to reproductive success. For decades, behavioral ecologists could not quantify reproductive success directly, and used proxies such as the number of observed mates or offspring produced (Emlen and Oring 1977; Klug 2011). Inferring reproductive success from such proxies involved making assumptions about species’ biology. For example, using the number of mates as a proxy for male fitness meant assuming that females do not vary in fecundity, and using the number of eggs in the nest of a breeding pair as a proxy for the male’s fitness meant assuming that the female does not engage in extra pair copulations or that occasional extra pair mates are unlikely to sire offspring.

However, in the last three decades, the advent of molecular means of assessing parentage has allowed direct and precise measurements of reproductive fitness, enabling novel insight into the complex landscapes of sexual selection acting both before and after copulation (e.g. Coltman et al. 2002; Birkhead 2010; Fisher and Hoekstra 2010). In many cases, these molecular measures have demonstrated that what we thought we knew about reproductive success was mistaken (e.g. Avise et al. 2002; Griffith et al. 2002; Uller and Olsson 2008; Boomsma et al. 2009). Specifically, biologists have discovered that the assumptions linking behavioral proxies to reproductive success were often not met. For example, females can vary in fecundity (Clutton-Brock 2009), may mate outside of observed social bonds (Griffith et al. 2002), and can store sperm, allowing for cryptic post copulatory female mate choice (reviewed in Eberhard 1996; Orr and Brennan 2015). In such cases, the reason for the mismatch between behavioral and genetic descriptions of mating systems is that, despite intensive field studies, researchers were yet to observe important components of a population’s mating system.

In this paper, we argue that mismatches between behavioral and genetic descriptions of mating systems can arise not only from undiscovered biology but also from the erratic and contingent progression of scientific research. In such a progression, poorly-supported conclusions from the earliest studies are inadvertently reified by later researchers, who, without examining the evidence for earlier conclusions, assume rather than test key hypotheses. Breaking away from such a progression of research is not inevitable, because it requires reinvestigating ideas believed to be true. Consequently, relatively unsupported corpora of knowledge about species’ social behavior and mating systems may remain undiagnosed.

We illustrate our argument using *Anolis* lizards, a model system for evolutionary ecology in which social behavior and mating systems have been studied for nearly a century (reviewed in Losos 2009). These decades of behavioral research yielded the near-unanimous conclusion that anoles are territorial and polygynous. In a chapter reviewing behavioral descriptions of *Anolis* mating systems, Losos (2009) concluded that “as a rule, male anoles are highly territorial.” Elsewhere, some of the best studied species in this genus have been described, based on behavioral observations, as matching “the paradigm of a territorial polygynous species” (Schoener and Schoener 1982). In what remains one of the best studies of anole social behavior in the wild, Rand (1967a), described their mating system thus:

> “…the lizards live together more or less permanently and the females usually mate with a single male (the male with the one or more females that have home ranges within his).”

Tokarz (1998), describing the prevailing views from behavioral data on anole mating systems, said that it is “generally believed that in territorial species of lizards, females that reside within a given male’s territory would have relatively few opportunities to mate with more than one male.” Stamps (1995) summarized their mating system as follows:

> “During the breeding season, male anoles defend territories that enclose the home ranges of adult females, and defend these mating territories against conspecific males. Although DNA paternity studies are not yet available for anoles, males probably father most of the hatchlings produced by the females within their territory.”

Together, these quotes help to delineate the prevailing view of anole spatial and social organization based on behavioral data. Under this view, which we describe as “strict territorial polygyny” and illustrate in Fig. 1, males have the potential to mate with one or more females within their territory, but females mate with only the one male in whose territory they are contained. If these territories are maintained for the duration of the breeding season or longer, as suggested by Rand (1967a), then all of a female’s offspring are expected to be sired by a single male.

**Figure 1.**
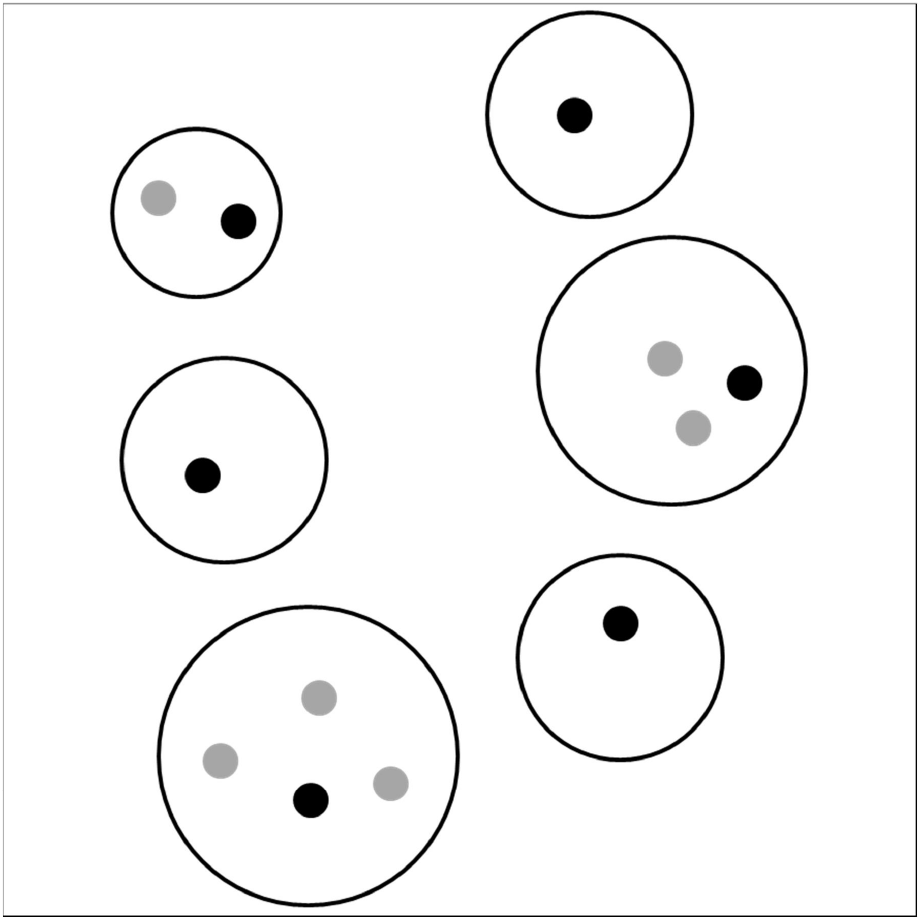
A pictorial representation of strict territorial polygyny. i.e. males (black) may mate with multiple females (grey) within their territories (black circles), but females mate with just the one male in whose territory they are contained. If this spatial organization is maintained for the duration of the breeding season, then all of a female’s offspring will be sired by just one male.

However, all the genetic evidence collected subsequent to these descriptions indicates that, as in many other reptiles and amphibians (Uller and Olsson 2008), females anoles’ offspring are frequently sired by multiple males; therefore, the prediction about strict territorial polygyny in *Anolis* lizards was not met (reviewed below; Passek 2002; Calsbeek et al. 2007; Johnson 2007; Harrison 2014). Quite to the contrary, female multiple mating is common in anoles, calling into question the behavioral descriptions predicting that female anoles will mate with just one male. Nevertheless, anoles continue to be described as territorial and polygynous (e.g. Calsbeek et al. 2007; Losos 2009; Simon 2011; Flanagan and Bevier 2014; Bush et al. 2016).

At the heart of this discrepancy between behavioral predictions and genetic data on female mating patterns in anoles is the concept of territoriality. Though territoriality is central to the behavioral descriptions of mating systems in many animals (Emlen and Oring 1977; Fitzpatrick and Wellington 1982; Lott 1984), the term itself is fraught with inconsistency and imprecision across different studies. Most often, the term “territorial” is used to describe individuals that defend an exclusive area in a fixed spatial location (Tinbergen 1957; Stamps 1977; Martins 1994; Maher and Lott 1995), indicating that the definition of territoriality incorporates two features: site fidelity (the tendency of an individual to remain in or return to a fixed spatial location) and exclusivity (the tendency of an individual to exclude other individuals, particularly conspecifics of the same sex, from the area they occupy). Under the strictest interpretation of territoriality in *Anolis* (Fig. 1), females mate with just one male; however, more relaxed interpretations of territoriality incorporating some variation in site fidelity, exclusivity, or both, can be consistent with female multiple mating. Imprecise and changing interpretations of territoriality across studies of anole social behavior may therefore have played an important role in producing the mismatch between behavioral and genetic descriptions of their mating system.

In this paper, we trace the evidence for territoriality, and for the relationship between territoriality and the expectation of polygynous mating patterns, in *Anolis* lizards. To this end, we examine nearly a century of research on their mating systems (see the Appendix for a list of papers considered). Our goal is to discern *how* we came to expect that female anoles mate with just one male when in fact they frequently mate with multiple males. Specifically, we examine if this research was somehow set on a path towards reifying a particular conception of territoriality that is inconsistent with widespread female multiple mating, leading to the erroneous expectation that anoles show strict territorial polygyny (Fig. 1). Throughout, we highlight whether the definitions and interpretations of territoriality employed by different researchers include site fidelity, exclusivity, or both; further, we pay attention to whether variation in site fidelity and exclusivity that could have explained female multiple mating remained undetected or was otherwise ignored.

We show that current ideas about anole social structure originated in studies whose scope and content is not commensurate with the weight they currently bear. These equivocal demonstrations of territorial behavior in early studies were seemingly taken at face value by later researchers, whose research included implicit and explicit assumptions about the existence of territoriality. Consequently, the design of later studies was often such that these studies were unable to detect variation in site fidelity and exclusivity. Moreover, even when later researchers found evidence for departures from strict territorial polygyny, this evidence was often deemphasized or ignored during data analysis and in the discussion of results. Given that mismatches between behavioral and genetic descriptions of mating systems are taxonomically widespread, our historical investigation reveals concerns that are likely not unique to *Anolis*. Indeed, the extent to which such erratic progressions of research afflict our understanding of animal behavior remains entirely unknown, and we urge researchers studying other organisms or questions to consider if the issues we highlight might apply to their fields of study as well. We conclude by considering the broader consequences of our case study for future research on animal mating systems.

## The Earliest Studies of Anole Social Interactions

The first study of lizard mating systems—Noble and Bradley (1933)—combined a review of existing natural history literature with laboratory observations on a taxonomically wide variety of lizard species. Both the lizards’ survival (“less than a year” for five species of *Anolis*, which typically live for at least a year even in the wild; Losos 2009) and their behavior indicated that the conditions under which these lizards were housed were likely stressful. Nearly half of all instances of copulatory behavior observed in *Anolis* by Noble and Bradley (1933) was between males. While this behavior was recognized as unusual, it was nonetheless interpreted as supporting territoriality—because lizards frequently engage in male-male copulations only in the lab, in nature these male-male copulations must be prevented by *something*.

This “something” was concluded to be the maintenance of exclusive territories, as evidenced by males’ propensity for aggression toward one another. Noble and Bradley (1933) remarked that “males tend to fight, and would, no doubt, tend to mark out territories for themselves.” Later, they said, about lizards in general, that “the only mechanism which is present to prevent males from copulating with other males as frequently as with females is that males when meeting each other during the breeding season tend to fight. The result is that males tend to occupy discrete territories, which are difficult to recognize in the laboratory but which have been described in the field.” The field studies of *Anolis* behavior referenced by Noble and Bradley (1933) only describe male-male aggression, and not site fidelity by either males or females. Thus, the existence of territoriality in anoles was first concluded on the basis of male-male aggression.

Evans (1936a, b, c) also concluded from laboratory experiments that male and female *Anolis* lizards maintain territories. Evans (1936a, c) detailed a weight-based social hierarchy among male *Anolis carolinensis* based on their aggressive interactions, which were described as the “urge to hold territory.” Again, conclusions were extrapolated from cages, in which animals were kept at high densities, to the field. For example, Evans (1936c) suggested, without reference to field data, that “the behavior of caged male *Anolis* is probably a modification of the behavior in the field. Under natural conditions when a strange male approached a particular territory which is in possession of another, a fight results…the beaten male retreats, leaving the victor in possession of the territory.”

Evans’ (1938a) subsequent field study was the first systematic research on anole territorial behavior in nature. Watching a population of *Anolis sagrei* for about a month, Evans (1938a) concluded that “*Anolis sagrei* exhibits a strong urge to select and defend a definite circumscribed territory.” Though this conclusion was largely based on observations of male-male aggression, Evans (1938a) also said that “proof that the species is territorial is given by the fact that the same individual has been observed many times on consecutive days upon a particular territory.” This dual approach indicates that Evans (1938a) included site fidelity as well as exclusivity in his conception of territoriality. Fortuitously, Evans (1938a) included transcriptions of all field notes taken during this study, which reveal that he concluded site fidelity based on a mean of three distinct observations per lizard. Though his systematic field-based approach was certainly path-breaking for its time, three observations made within a short period relative to the full breeding season (*A. sagrei* breed for at least six months; Tokarz et al. 1998) cannot be considered sufficient to demonstrate persistent site fidelity.

Critique from Evans (1938a, b) prompted Greenberg and Noble (1944) to modify the conditions under which observations were conducted in the lab—they housed and observed *A. carolinensis* lizards in larger cages and greenhouses, up to 5 m ×5 m. But these larger arenas may still have been too small to assess if the multiple males they contained each maintained exclusive areas and showed site fidelity. The authors mentioned that “an active adult male usually succeeded in dominating the entire cage,” which implies that males in these cages did not maintain exclusive areas, potentially an artefact of a small arena size. The conditions in the cage were nonetheless described as “near-normal competitive conditions.”

Oliver’s (1948) methods for observing *A. sagrei* in the Bahamas were similar to Evans’ (1938a)—17 lizards in an area approximately 4 × 20 m were “marked and casually observed for a period of slightly less than one month.” And though Oliver (1948) “planned to present elsewhere at a later date a detailed account of the individual and social activity of this species,” to the best of our knowledge, no such account was published. Oliver (1948) summarized his results as showing that “definite territories are maintained and defended by both sexes.” However, the territories he described were not exclusive, because “within the area occupied by each large male there was a smaller male,” and it is not clear if these smaller males were reproductively active or not. His conception of territoriality in anoles was therefore potentially consistent with female multiple mating.

Approximately contemporaneous natural history studies described anoles as territorial based on far less evidence. For example, Thompson (1954) observed a single male *A. carolinensis* displaying at a “jar containing about a dozen swifts (*Sceloporus undulatus*) that I had collected the day before,” as well as at a skink, and concluded that “during the entire performance it seemed that the anolis [sic] might have been trying to hold or establish a territory.” In sum, these early studies of anole social behavior all readily described these lizards as territorial, despite presenting limited data that were insufficient to demonstrate site fidelity and did not always demonstrate exclusivity.

## The Firm Establishment of Territorial Polygyny

In the decades that followed these early studies, territoriality remained a frequently used description for anole space use behavior and social interactions; the next watershed moments in this research trajectory came when these descriptions grew to explicitly include a polygynous mating system.

In what remains one of the most detailed studies of *Anolis* territoriality, A. Stanley Rand spent almost a year observing the movement patterns and social interactions of *Anolis lineatopus* in Jamaica. This yielded a paper in which Rand (1967a) fully expressed the tension between adhering to a territorial framework on one hand, and observing variation in site fidelity and exclusivity on the other. Nonetheless, Rand (1967a, b) proposed a tight link between territoriality and polygyny based on the idea that males maintain exclusive mating access to females.

At least part of Rand’s (1967a) conception of territoriality was derived from earlier research on anoles. For example, he cited Evans (1938a) in describing the pattern of “a male with a home range shared by one or several females that are his mates” in *A. sagrei*. He also suggested that *A. lineatopus* and *A. sagrei* have similar social behavior based on Oliver’s (1948) description of the latter as territorial. But Rand (1967a) also demonstrated the complications of fitting messy field data into this territorial framework.

These complications are best captured by Rand’s (1967a) descriptions of these lizards’site fidelity. First, he stated that “an *A. lineatopus* seldom travels far and most of the area it visits is visible to it from its usual perch.” But following this he describes how, in calculating the area over which an individual lizard is active, he “omitted the occasional visits that certain *A. lineatopus* made to perches well outside of the area where they were usually seen.” Thus departures from site fidelity that may have been reproductively important were excluded while attempting to establish site fidelity.

A similar dissonance was also evident when Rand (1967a) first stated that “the activity range of an adult *A. lineatopus* seems relatively permanent and certainly shows no seasonal variation” but then described data that may have suggested otherwise. Documenting the locations of 16 adult males in one of his field sites, he noted that these males were seen multiple times while sampling in September and October but only seven of these—less than half—were still present in the site five months later. Rand (1967a) acknowledged that “of those nine which had not been seen in March, two were dead, but it is possible that the other seven had shifted their areas outside of the study plot.” In other words, Rand (1967a) considered that almost half of the adult males in this site may have shown seasonal departures from site fidelity, but nevertheless concluded that these lizards remain in fixed locations permanently.

Rand’s (1967a) thoughts on exclusivity were complex, illustrated by his statement that “individual aggression may be expressed as either of two types: dominance hierarchies and territoriality…The behavior of *A. lineatopus* can not be assigned to either of these categories because it has important aspects of each of them.” He went on to explain that while “every *A. lineatopus* holds a territory, defending it against neighbors of the same size…each is a member of a straight line dominance hierarchy that consists of all those anoles of different sizes whose home ranges overlap its own home range.” Because large as well as small males were observed mating, such a spatial organization appears inconsistent with the idea that males maintain exclusive mating access to the females within their territory.

Despite these dissonances and complexities, Rand (1967a) unequivocally linked territoriality to polygyny, by proposing that male territoriality is adaptive in *Anolis* because it allows males to maintain exclusive mating access to females:

> “I think the general occurrence of aggressive behavior and the spacing out it produces in all sizes of *A. lineatopus* can be explained by…ecological advantages…but the greater aggressiveness of the adult males requires additional explanation. I think the explanation lies in a function of territory discussed at length by Tinbergen (1957), which demonstrates the selective advantage that is conferred on an adult male if he can insure himself exclusive mating rights to certain females by keeping other males away from them. If he can do this for a single female, he insures that he will father at least some offspring, and the more females he can keep isolated, the more offspring he will have and the greater his contribution to the gene pool of the next generation. This being true, there must be a strong selection pressure for any mechanism that will insure a male exclusive mating rights to one or more females. The aggressive behavior of adult male *A. lineatopus* that keeps other males out of the area in which females are permanently living is just such a mechanism.”

In a second paper based on these data, Rand (1967b) again concluded that while all individuals defend territories for access to food, males also defend access to mates, thereby reinforcing the link between territoriality and polygyny in *Anolis*. This idea that males maintain exclusive mating access to females was almost certainly a sign of the times. Hinde (1956), in his introduction to an issue of *Ibis* devoted to territoriality in birds, proposed a hypothesis similar to the one espoused by Rand (1967a, b): “Any behaviour of the male which helps to prevent his mate being fertilized by another male is likely to carry a great selective advantage.” This notion of the “monopolizability” of females, or of the resources to which females are attracted, became the foundation of how behavioral ecologists understand the evolution of animal mating systems (Orians 1969; Emlen and Oring 1977). In anoles, it was quite possibly the basis of the expectation of strict territorial polygyny, which rests on the assumption that males maintain exclusive mating access to the females in their territory (Fig. 1).

Though research on anole mating systems grew rapidly after 1967 (discussed below), the next major step towards firmly establishing the link between territoriality and polygyny came 17 years later. Ruby (1984) examined male breeding success in *A. carolinensis* in the context of space use, motivated by the assessment that “mating systems of reptiles are poorly known…and formative factors remain undetermined.” Sampling for over five months for each of two consecutive years, including daily observations for three months each breeding season (though over only a 460 m2 area), Ruby (1984) discovered ways in which these lizards’ behavior did not conform to the expectations of territorial polygyny that were laid out by Rand (1967a, b). For example, he noted that “only 17 of the 68 (25%) males remained 12 weeks or longer during a single breeding season of 20 weeks,” potentially indicating variation among males in site fidelity. Moreover, he found that “female [territories] overlapped more than one male in about 25% of the receptive periods [two week intervals in the breeding season]” and in calculating the number of potential mates of males, each “female was assigned to all overlapping males.”

These observations and analytic choices indicate that Ruby (1984) uncovered the potential for females to mate with multiple males, and thus documented a mating system in which males do not maintain exclusive mating access to individual females. Ruby (1984) even considered the possibility that sperm storage is an adaptation for female mate choice in these lizards. Nonetheless, at the very outset of the paper, Ruby (1984) proposed that mating systems in lizards range from monogamy to polygyny and described territoriality as “one means of gaining exclusive mating access to females.” Later in the paper, he stated that “because the *Anolis* breeding system appears to be resource defense polygyny (Emlen and Oring 1977), territoriality is favored as a means of restricting access to mates.” It is possible that Ruby’s (1984) data led him to soften his stand from expecting males to maintain “exclusive” mating access to expecting “restrict[ed]” mating access; nonetheless, Ruby (1984) was subsequently frequently cited as supporting the idea that anoles are territorial and polygynous without explicitly acknowledging this potential for female multiple mating (e.g. Qualls and Jaeger 1991; Stamps 1995; Jenssen et al. 2000, 2005; Lovern 2000).

## The Consequences of Limited Sampling

Research on anole behavior blossomed between Rand (1967a, b) and Ruby (1984). However, because by this point the consensus seemed to be that anoles are territorial, this research was not often designed to explicitly test if these lizards behave territorially, i.e. to demonstrate that they exhibit site fidelity and exclusivity. Specifically, territoriality was an almost foregone conclusion in studies with a limited spatial and temporal extent of sampling. In other words, the design of many of these studies was such that they were unlikely to uncover evidence that individual anoles vary in site fidelity or exclusivity, and therefore were unlikely to point to the possibility that females often mate with multiple males

If the sampling period of a study of social behavior is not long enough, then relatively infrequent but reproductively consequential departures from either male-male exclusivity or site fidelity may not be detected often enough that they are considered signal and not noise. For site fidelity, this includes not only occasional forays away from and returns to a fixed territory, but also shifts in territory location that may take place only a few times per breeding season—neither would be detected by studies with short durations. An extreme example of a constrained sampling period can be seen in Philibosian’s (1975) study of *Anolis acutus* and *Anolis cristatellus*, in which he stated that “often an observation period of one day was sufficient to record enough positions and enough encounters involving the residents on a tree to make reasonably accurate territory descriptions.” As researchers became more certain that anoles are territorial, they became comfortable making more extreme assumptions. For example, in estimating the number of neighbors of individual *A. sagrei*, Calsbeek (2009) estimated the center of a lizard’s territory as simply the first location at which that lizard was observed.

Moreover, if a study of social behavior does not sample over a large enough area and a sampled individual disappears from the study site, researchers cannot know if the individual has died or simply moved. Thus, studies with limited sampling areas will be most likely to sample only those individuals who stay in the same place. For example, Trivers (1976), studying the Jamaican *Anolis garmani*, “attempted to map male territories by concentrating on a small portion of the study area.” He stated that “males are sighted too infrequently to measure territory size the usual way; that is, to construct a volume fitting such sightings.” These infrequent sightings could conceivably be due to the low chance of re-spotting individuals with low site fidelity within a small area. But Trivers (1976) continued by saying that “fortunately males 105 mm and larger show a strong tendency to occupy trees…Typically, during a given visit, a large male will be sighted between five and ten times in a large tree.” Thus, Trivers (1976) focused his sampling for estimating territory size to a small area known to be occupied by individuals with high site fidelity, limiting the variation in movement behavior that could be detected.

The combination of spatially and temporally restricted sampling can be seen in work by Jenssen and colleagues (e.g., Jenssen et al. 1995; Jenssen and Nunez 1998), who documented the behavior of a population of *A. carolinensis* along the Augusta Canal in Georgia. This population inhabited a thin strip of vegetation (three to six meters wide), which comprised clumps of trees observable from an elevated walkway, and the activity of lizards in each clump of trees was watched for only eight days, out of a months-long breeding season. Nonetheless, these data were interpreted to conclude that “males are polygynous, defend closely monitored and stable territories, and devoted large blocks of time and energy on territory maintenance” (Jenssen et al. 1995). With time, statements of territorial polygyny thought to be supported by these data became even stronger, such as this statement from Jenssen et al. (2000): “the *A. carolinensis* mating system is driven by the outcome of intermale territorial aggression. Winners achieve and maintain direct mating access to varying numbers of females…because females are relatively sedentary and clustered in small contiguous home ranges.”

It is certainly worth noting that while the sampling design in these studies reveals, with hindsight, certain assumptions regarding territoriality, Jenssen and colleagues’ fieldwork simultaneously challenged other beliefs that were commonly held by laboratory-based researchers studying anole behavior. For example, using similar sampling methods to those described above, Jenssen et al. (2001) tested and found no evidence for the hypothesis, long held by neuroendocrinologists, that male *A. carolinensis* emerge at the end of the winter and establish territories prior to female emergence.

## Four Fates of Documented Departures from Territoriality

Evidence for variation in territorial behavior, namely the extent of site fidelity and exclusivity, was implicitly and explicitly excluded through much of the later literature on *Anolis* social behavior. This exclusion took on at least four different forms. The first and second forms correspond to what is known as the “primary simplification” of scientific research, whereby the construction of facts is influenced by scientists’ decisions on how to present the data in a paper (Dewsbury 1998).

In the first form, already seen in Rand (1967a), departures from territoriality were removed at the time of analysis. For example, Trivers (1976) quantified male *A. garmani* territory sizes based on the size of trees that individuals occupied, and “a tree was assigned to a male if he was seen three or more times in it without any other adult male being seen therein.” However, “if, as happened several times, a large tree was also known to be occupied by a small adult male (85 mm – 104 mm), both males were excluded from the data, since too few data were available to partition the tree between them,” even though male *A. garmani* as small as 87 mm in size were observed copulating with females. Thus, departures from male-male exclusivity were explicitly excluded when considering these lizards’ territoriality. Similar choices were also made in considerations of site fidelity. For example, Schoener (1981) argued that in calculating home range areas based on location data, “the inclusion of the outermost observations…may still be undesirable” because “the utilization may resemble a more compact distribution if outliers were disregarded.” As a result, the home ranges of four anole species in the Bahamas were calculated without including the “10% of points farthest from the geometric center” (Schoener and Schoener 1982). While this analytic choice is certainly justifiable for calculating the centers of individuals’ activity, it compromises the ability to predict mating patterns from space use behavior, unless one is certain that individuals do not mate when at the 10% of points farthest from the geometric center.

A second fate of observed departures from territoriality, as seen in Ruby (1984), involved quantifying them but omitting them from interpretation. For instance, Schoener and Schoener (1980) describe *Anolis sagrei* as exemplifying the “paradigm of a territorial, polygynous species” even though between 3% and 28% of males in six populations remained within their study sites for less than a week, potentially indicating frequent deviations from site fidelity. An implicit justification for ignoring this often substantial proportion of males from a description of the lizards’ mating system is that these “floating” males do not mate with females. Though this is a reasonable and testable hypothesis, *assuming* that non-territorial males do not reproduce simply because they are not territorial is unjustified. In another example, Fleishman (1988) categorized adult male *Anolis auratus* as either territorial or non-territorial, based on their display behavior and levels of aggression. Even though non-territorial males were observed copulating with females within the territories of territorial males, Fleishman (1988) stated that “territories of *Anolis* males are primarily for exclusive access to mates.”

In a third, distinct fate, research that explicitly documented departures from territoriality stayed unpublished and had little influence. Consider two abstracts submitted to the annual meeting of the Society for Integrative and Comparative Biology. Both studies (Alworth 1986; Webster and Greenberg 1988) examined *A. carolinensis* behavior in enclosures. While Webster and Greenberg (1988) found that “the average site fidelity was 52%,” Alworth (1986) concluded that “territoriality in these lizards [should] be regarded as a highly flexible behavioral tactic adaptive only in specific contexts” and that “the broad characterization of a genus or species as territorial is misleading.” However, to the best of our knowledge, neither of these studies was published.

Finally, in the fourth fate, deviations from territorial polygyny in *Anolis* were documented and acknowledged fully, but the species’social behavior was described as an exception to the rule. For example, *Anolis valencienni* was described by Hicks and Trivers (1983) as displaying “many features atypical of other *Anolis*,” including the lack of territorial behavior by either males or females. Consequently, “because many adults of both sexes encounter each other daily, there are unusual opportunities for female choice…over a period of six weeks, a female may copulate with five or more males.” This “unusual” opportunity for female multiple mating was hypothesized to be due to *A. valencienni’s* tendency to forage more actively than other anoles. We are not suggesting that *A. valencienni* does not differ in its behavior from other anoles; in fact, its behavior must be different enough that it was recognized as exceptional by researchers working within the paradigm of territorial polygyny. But because *A. valencienni* was positioned as exceptional, its behavior was never cause to re-evaluate the behavior of other anole species.

## Two Exceptions

In seven decades of research on anoles, two studies explicitly described these lizards’ social behavior as being consistent with female multiple mating. The first—Gordon (1956)—remained relatively uninfluential, but the second—Tokarz (1998)—laid the groundwork for the reconciliation of behavioral observations with subsequent genetic studies that in fact detected evidence for female multiple mating.

In his dissertation, Gordon (1956) aimed “to analyze, biodemographically, two local populations” of *A. carolinensis*. The work comprised primarily of nocturnal censuses in two 20 m × 20 m plots every two weeks for over a year, with all captured individuals marked permanently. Gordon’s (1956) data revealed the potential for departures from site fidelity: 73% of 1024 marked lizards were observed just once within the study site, and only 8% of all lizards, and 13% of adults, were observed three or more times. Though some of the disappearances were undoubtedly due to predation and others must have resulted from the failure to detect individuals again, the data are also consistent with many individuals in this population exhibiting low site fidelity. Gordon (1956) later questioned anoles’ site fidelity when describing lengthy disappearances of individual lizards from the study site and frequent long distant movements. He also wrote the following:

> “The individual female may copulate with more than one male per season. The social group is maintained by the activity of the dominant male, and sexual bonds between the male and his females are loosely formed. Females tend to wander more than males and ample opportunity is present for a female to be attracted to, and take up residence in, another male’s territory. In cases of territorial hierarchy, the dominant male and his subordinates may share the same group of females.”

Though it certainly had the potential to do so, Gordon’s (1956) thesis did not end up provoking a shift in how behavioral ecologists think about anole mating systems. For example, three influential papers on *Anolis* territorial behavior (Schoener and Schoener 1982; Ruby 1984; Jenssen et al. 1995) cite Gordon (1956) but do not refer to his suggestion that female anoles may readily mate with multiple males.

Over four decades later, behavioral observations by Tokarz (1998) demonstrated even more clearly that female *A. sagrei* have the opportunity to mate with multiple males. He explicitly questioned the idea that males maintain exclusive mating access to females in their territories, saying that “few studies have attempted to record the mating pattern of individual females in nature as a means of evaluating the potential for female mate choice and sperm competition.” Tokarz’s (1998) data revealed that “most females (75%) had more than one mating partner, and this was due almost entirely to females mating with new males that successfully supplanted previous males from their territories.” A decade later, however, Tokarz (2008) minimized his own previous findings, saying that “male territories in *A. sagrei* appear to be relatively stable at least during the midsummer portion of the breeding season (Evans, 1938[a]), although instances of males being supplanted from their territories by other males have been observed (Tokarz, 1998).”

It is tempting to conclude that Tokarz’s (1998) results solve the problem of the mismatch between behavioral and genetic descriptions of anoles’ mating system. To an extent, they do, but his documentation of turnover in male territory occupancy is only one of many different ways in which departures from strict territorial polygyny (Fig. 1) could facilitate female multiple mating. Other ways, such as multiple reproductive males occupying overlapping areas, had been documented in anoles by previous researchers, but their potential relevance to female multiple mating was downplayed. Yet other ways, such as the existence of reproductive males or females who wander non-territorially, are unlikely to be detected in studies with small sampling areas or durations. This includes Tokarz’s (1998) study, in which 16 individuals occupying a single tree that was 2 m in diameter, were watched for just over a month. That said, even Tokarz (1998) observed “six instances in which males…entered an adjoining male’s territory and courted females there.”

These different possible routes to multiple female mating have different implications for anoles’ reproductive dynamics and sexual selection. Multiple mating resulting from male territorial turnover may lead to serial polygyny, in which at any one time, a territorial male is the exclusive mate of females residing within his territory. Alternatively, other types of departures from site fidelity and exclusivity lead to situations in which, at any given time, females may be able to mate with several males, allowing for female mate choice. While the serial territorial polygyny that Tokarz (1998) observed may certainly be a male adaptation for achieving high reproductive success, we cannot know from existing behavioral data if it is the only reproductive strategy, or even the dominant reproductive strategy, adopted by male anoles.

Crucially, it is not necessary that every individual in a population depart from site fidelity or exclusivity in the same way or to the same extent for the link between territoriality and polygyny to be compromised. There is therefore a disconnect across levels of biological organization that is central to reconciling behavioral and genetic descriptions of mating systems—while behavioral descriptions apply to individuals, the mating system is a population-level trait. Equally, different populations and species may also vary in the composition of reproductive strategies across individuals (Lott 1984; Kappeler et al. 2013), and the proportion of individuals in a population who behave territorially influences our ability to predict whether the population’s mating system will in fact be polygynous. This explanation also makes clear that many previous studies of anole social behavior that concluded that anoles are territorial may have accurately described the behavior of *some* individuals. However, to the extent that the results of existing genetic studies are general, previous behavioral studies either did not accurately describe the behavior of *all* individuals, or erroneously failed to consider as reproductively important those individuals whose behavior they described as deviating from territoriality. The disconnect between behavioral and genetic descriptions of a population’s mating system thus becomes quantifiable by considering variation across reproductive individuals in the extent to which their behavior differs from territoriality.

## The Age of Genetics

The use of genetic tools uncovered female multiple mating in three species of anoles—*A. carolinensis*, *A. sagrei*, and *A. cristatellus*. Each of these studies (one paper published in a peer reviewed journal, as well as three theses that, at present, are unpublished) discussed the implications of their findings for territoriality to different extents.

Passek (2002) examined the possibility for sperm choice or competition in *A. carolinensis* using a combination of behavioral and genetic approaches. She invoked variation in site fidelity and exclusivity when saying that “while males defend territories that contain multiple female home ranges (Jenssen et al. 1995), the potential exists for extra-pair paternity due to temporary invasion by “floater” males or female home ranges being overlapped by more than one male (Ruby 1984).” Though Passek’s (2002) description suggests only occasional departures from territoriality, her genetic data showed that 48% of offspring were sired by males other than the one identified as the territory owner, including 21% sired by smaller males within the same territory and 15% sired by neighboring males. The paternity of the remaining 12% of offspring could not be determined. In her conclusion, Passek (2002) expressed skepticism that anyone had accurately measured “the frequency of territorial exchanges resulting from territory takeovers.”

Johnson (2007) mapped *A. cristatellus* space use behavior over a three week period, and found that females’ “territories overlapped an average of 3.3 males.” Genetic data confirmed this potential for females to mate multiply, showing that “52% of females laid eggs sired by multiple males.” Moreover, variation in site fidelity also played a role in facilitating female multiple mating, because “26% of offspring were sired by males whose territories did not overlap that of the mother.” She concluded that “these results may be explained by a combination of a male dominance hierarchy…and female mate choice,” mating strategies and interactions that are not encompassed by strict territorial polygyny (Fig. 1).

In the only published evidence for multiple mating by female anoles, Calsbeek et al. (2007) found that “more than 80% of field-caught *A. sagrei* females that produced two or more progeny had mated with multiple males [making] *A. sagrei* one of the most promiscuous amniote vertebrates studied to date.” However, this paper did not tackle the implications of its results for territoriality.

Finally, the most direct evidence for departures from territoriality influencing anole mating systems again combined behavioral observations with genetic data (Harrison 2014). Studying *A. carolinensis*, Harrison (2014) assumed site fidelity in her behavioral sampling by mapping the home ranges of lizards after observing individual’s spatial locations for 30-minute focal observations (it is not clear how many focal observations were conducted for each individual; Harrison [2014] does mention that “behavioral observations were conducted at irregular intervals, making it difficult to determine whether males shifted their territories during the study period”). However, her genetic data revealed that spatial proximity, as determined by the focal observations, did not predict mating between pairs of males and females. In fact, the mean distance (± standard deviation) between mating pairs was 33±22 m, over five times the mean estimated territory diameter in that population. This indicates that individual lizards *must have* moved between when they mated and when they were observed. In the face of this evidence, Harrison (2014) continued to invoke a territorial paradigm to understand anole social behavior, at least initially: “males and females from opposite sides of the study site mated relatively frequently…often traversing distances over 60 m. For this to occur, either the male or female (or both) left its territory at some point, or they mated before establishing territories and used stored sperm.” Later, however, she proposed a number of hypotheses for male movement behavior, including the existence of an alternative non-territorial, wandering male strategy adopted by adult males, and temporal variation in individual site fidelity within a single breeding season, that definitely break out of the mold of territoriality.

## Broader Implications for Animal Mating Systems

This century-long trajectory of research on *Anolis* mating systems exemplifies several larger issues that could plague the study of animal mating systems more generally. However, it is challenging to establish that the problems we identify here are generally applicable, because discerning their applicability to a particular taxon demands a close familiarity with the full body of literature on that taxon’s biology, as well as familiarity with the organism’s biology itself. In this final section, we identify the main driving forces that led to the incomplete and possibly incorrect descriptions of *Anolis* social behavior, culminating in the erroneous prediction that each female’s offspring will be sired by the single male in whose territory she resides. We hope this discussion will prompt researchers who are intimately familiar with other organisms’ biology to re-examine the basis of what we think we know to be true about those organisms’ social behavior.

The history of research on *Anolis* mating systems demonstrates multiple ways in which the erratic and contingent progress of research may have prevented researchers from fully describing the behaviors that facilitate female multiple mating in these lizards. The central problem was described well by Stamps (1994), although she was discussing specific aspects of territoriality not covered in this review:

> “Current ideas about the behavior of territorial animals are based on a series of assumptions…in some cases these assumptions have not been adequately tested. By virtue of repetition, untested assumptions have a tendency to solidify into “quasi-facts.””

Such repetition certainly characterized the earliest studies of *Anolis* social behavior, where studies repeatedly concluded that anoles are territorial based on often flimsy evidence. It is not clear whether the authors of these earliest studies considered the implications of these lizards’ space use and movement patterns for their mating system. It is possible that territoriality was so readily assumed and concluded in these early studies *precisely because*, under the strictest interpretation, territoriality is incompatible with female multiple mating. Charles Darwin, in his seminal text on sexual selection, expressed the prevailing view at the time that females are generally “coy,” “passive,” and “less eager” to mate than are males (Darwin 1871; discussed in Hrdy 1986; Dewsbury 2005; Tang-Martinez and Ryder 2005; Tang-Martinez 2016). Moreover, many biologists at the time believed that females of most species were unlikely to possess the cognitive ability to make choices about which males to mate with, and ignored evidence to the contrary (reviewed in Milam 2010). Invoking a mating system such as territorial polygyny, which under the strictest interpretation leaves females unable to choose between males and assumes that females have no reason to seek out multiple mates, thus may have been a sign of the times.

However, Greenberg and Noble (1944) conducted experiments explicitly to test whether female anoles choose mates on the basis of males’ dewlaps, asking if females preferred to mate with males with intact or manipulated dewlaps. They found no effect of dewlap manipulation on mating success, but by asking the question, these authors revealed that they considered female mate choice possible in anoles, and thus considered that females have the opportunity to mate with multiple males. In contrast, later researchers studying anole territorial behavior frequently maintained that female mate choice was unlikely because it is precluded by territoriality. For example, Schoener and Schoener (1980) suggested that “adult females seem quite sedentary in [*A. sagrei*], and the opportunity for female choice would seem correspondingly limited,” and Stamps (1983), in a review of lizard territoriality and polygyny, said the following:

> “In most insectivores, female choice of mating partner is probably fairly limited. Since females do not leave their home ranges in order to mate, prospective male partners must have home ranges overlapping that of the female. A female with a home range on the border between 2 male home ranges might be able to choose between them, but this option is restricted in territorial species by the males’ tendencies to arrange their territories to completely enclose female home ranges.”

Thus, though researchers all the way from Noble and Bradley (1933) to Stamps (1983) and beyond described anoles as territorial, the predictions for mating patterns derived from that behavioral description, such as whether females have the opportunity to choose mates, could be inconsistent with one another.

That the term “territoriality” as interpreted by different researchers could be compatible with fundamentally different expectations for patterns of mating and sexual selection highlights the fact that very few studies define territoriality explicitly (Maher and Lott 1995). Different authors’ conceptions of territoriality include different degrees of variation in both site fidelity and exclusivity, and therefore lead to different expectations for female multiple mating. This fuzziness in the definition of territoriality also raises the following question—at what point might we conclude that territoriality is too imprecise a term to be useful as a predictor of a species’ mating patterns? Departures from male-male exclusivity have been observed in anoles (e.g. Rand 1967a; Trivers 1976; Fleishman 1988), but these examples were still considered to be within the fold of territoriality because “exclusivity” was qualified or limited to mean that males only exclude size-matched individuals. These qualifications were made even though males in smaller size categories were observed to mate with females. Similarly, a lack of clarity about the meaning of site fidelity permeates research on territorial behavior—does “site fidelity” mean staying in the same place, leaving but always returning to the same place, or attempting (but possibly failing) to stay in or return to the same place? How long does an individual have to stay in a certain place to be considered site faithful? Almost all possible answers to these questions have, at some point in the last century, been implicitly or explicitly accepted as consistent with territorial behavior in anoles, even though each answer can lead to very different expectations for mating patterns.

Once territoriality became established as a description of anoles’ mating system, the design and interpretation of subsequent studies of these lizards’ social behavior made it difficult to detect variation among individuals in site fidelity or exclusivity, variation that could easily be reproductively consequential. Which individuals were studied, the extent of sampling area and duration, the data that were analyzed versus excluded, and the extent to which inconsistent findings were deemphasized—each of these scientific decisions involved choices that would determine whether the study could actually test the precepts of territoriality or whether it simply assumed them. For the most part, the choices made were such that territoriality remained untested. However, these studies were written and interpreted as if the idea that anoles are territorial had been tested, and thus each seemed to provide independent confirmation of this description of their spatial and social organization. In fact, even though these studies were conducted by different researchers on different populations and species of anoles, they were conceptually non-independent, unintentionally leading the earliest studies to “assume a stature that their original authors never intended” (Stamps 1994).

It is this problem—adhering to a conceptual paradigm while designing studies that are consequently unlikely to uncover or take seriously the evidence that would allow you to escape that paradigm— that we believe is the most important problem revealed by our review. This problem cannot be solved simply by collecting more data; reaching a solution additionally requires that we explicitly identify and question the assumptions made when designing research (Gowaty 2003). But framing the challenge thus also makes the solution clear—we should continue collecting observations of animals’ behavior in a manner that is as free as possible from existing conceptual frameworks, even in taxa whose biology we think we know well. In other words, the solution calls for renewed and continued attention to organisms’ natural history (Greene 2005; Tewkesbury et al. 2014). As Greene (2005), who defined natural history as “descriptive ecology and ethology,” put it, “discoveries of new organisms and new facts about organisms often reset the research cycles of hypothesis testing and theory refinement that underlie good progressive science.”

The call for a close relationship between natural history observations and the advance of research in animal mating systems is far from new. We conclude with a remarkably apt excerpt from a 1958 letter to the editor of *Ibis* from John T. Emlen, following an issue about territoriality in birds (Hinde 1956):

> “There is a growing tendency among ornithologists to blindly and devotedly follow what is becoming a fixed or conventional concept of territory. Instead of describing their observations directly, authors often seem to go out of their way to fit them into the “accepted” pattern through the “approved” terms and phrases.”

Emlen (1958) continued:

> “My concern in this letter is with the tyranny of words and with the dangers inherent in patterned thinking. The fascination of catch phrases and the reverence with which they come to be held are major, though subtle, obstructions to free and accurate thinking. Conventionalized phrasing, furthermore, often leads to conventionalized thinking, the very antithesis of free investigation and the arch-enemy of scientific progress. A neat, substantive definition of territory has the fascination of finality, but in a virile science dead ends must be avoided, not sought; it has the fascination of authority, but basically we recognize that the study of natural phenomena must not be subordinated to the study of intellectual creations.”

The accurate quantification by genetic means of individuals’ reproductive success in natural populations is valuable not just because such data help to render more complete descriptions of animals’ social and reproductive behavior. These data also let us identify taxa in which the erratic and contingent progression of scientific research may have led behavioral ecologists towards erroneous conclusions about animals’ mating systems. But the genetic data alone do not shed light on the question of how we come to believe such conclusions. We contend that taxon-specific historical investigations into this question allow us to escape the confines of “conventionalized phrasing” and “conventionalized thinking,” and are an important step towards designing studies that will let us understand animal social behavior in its full complexity.

## Acknowledgments

Stamps and R.R. Tokarz gave us valuable feedback on previous drafts that substantially improved the manuscript, as did C.M. Donihue, Y.E. Stuart, S.R. Prado-Irwin, P. Muralidhar, M.E. Kemp, N.E. Herrmann, M.R. Lambert, E.E. Burnell, D.P. Rice, J.H. Boyle, L.J. Martin, and two anonymous reviewers.

## Appendix: Papers examined

A list of all the papers examined in our historical investigation of territorial polygyny in Anolis lizards, in alphabetical order. We searched for papers on Web of Science using keywords “Anolis” or “Norops” and “territor*”. From the results, we selected papers that were directly relevant to Anolis territoriality, in that the authors studied male-male aggression or site fidelity, including mapping home ranges, or based their study or discussion of Anolis social or reproductive behavior on prior conclusions of territoriality. We also followed relevant citations from within the sampled papers, yielding a set of 106 papers that spanned over nine decades and included field- and lab-based studies, as well as conceptual papers and reviews.

## REFERENCES

Alworth TJ (1986) Perch availability and season affect aggression levels in the territorial lizard, Anolis carolinensis. Am Zool 26:1041

Avise JC, Jones AG, Walker D et al (2002) Genetic mating systems and reproductive natural histories of fishes: Lessons for Ecology and Evolution. Annu Rev Genet 36:19–45

Birkhead TR (2010) How stupid not to have thought of that: post-copulatory sexual selection.

J Boomsma, Kronauer DJC, Pedersen JS (2009). The evolution of social insect mating systems. In: Gadaue J, Fretwell J (eds) Organization of Insect Societies. Harvard University Press, Cambridge, pp 3–25

Bush, JM, Quinn MM, Balreira EC, Johnson MA (2016) How do lizards determine dominance? Applying ranking algorithms to animal social behaviour. Anim Behav 118:65–74

Calsbeek R (2009) Sex-specific adult dispersal and its selective consequences in the brown anole, Anolis sagrei. J Anim Ecol 78:617–624

Calsbeek R, Bonneaud C, Prabhu S, Manoukis N, Smith TB (2007) Multiple paternity and sperm storage lead to increased genetic diversity in Anolis lizards. Evol Ecol Res 9:495–503

Clutton-Brock T (2009) Sexual selection in females. Anim Behav 77:3–11

Coltman DW, Festa-Bianchet M, Jorgensen JT, Strobeck C (2002) Age-dependent sexual selection in bighorn rams. Proc R Soc Lond B 269:165–172

Darwin C (1871) The Descent of Man, and Selection in Relation to Sex. John Murray, London

Dewsbury DA (1998) Robert Yerkes, sex research, and the problem of data simplification. Hist Psychol 1:116–129

Dewsbury DA (2005) The Darwin-Bateman paradigm in historical context. Integr Comp Biol 45:831–837

Emlen JT (1958) Defended area?—A critique of the territory concept and of conventional thinking. Ibis 99:352

Emlen ST, Oring LW (1977) Ecology, sexual selection, and the evolution of mating systems. Science 197:215–223

Evans LT (1936a) Social behavior of the normal and castrated lizard, Anolis carolinensis. Science 83:104

Evans LT (1936b) Territorial behavior of normal and castrated females of Anolis carolinensis. Pedagog

Semin J Genet Psychol 49:49–60

Evans LT (1936c) A study of a social hierarchy in the lizard Anolis carolinensis. Pedagog Semin J Genet Psychol 48:88–111

Evans LT (1938a) Cuban field studies on the territoriality of the lizard Anolis sagrei. J Comp Psychol 25:97–125

Evans LT (1938b) Courtship behavior and sexual selection of Anolis. J Comp Zool 26:475–497

Fisher HS, Hoekstra HE (2010) Competition drives cooperation among closely related sperm of deer mice. Nature 463:801–803

Fitzpatrick SM, Wellington WG (1982) Insect territoriality. Can J Zool 61:471–486

Flanagan SP, Bevier CR (2014) Do male activity level and territory quality affect female association time in the brown anole, Anolis sagrei? Ethology 120:365–374

Fleishman LJ (1988) The social behavior of Anolis auratus, a grass anole from Panama. J Herpetol 22:13–23

Gordon RE (1956) The biology and biodemography of Anolis carolinensis carolinensis Voight. Dissertation, Tulane University

Gowaty PA (2003) Sexual natures: How feminism changed evolutionary biology. Signs 28:901–921

Greenberg B, Noble GK (1944) Social behavior of the American chameleon (Anolis carolinensis Voight). Physiol Zool 17:392–439

Greene HW (2005) Organisms in nature as a central focus for biology. Trends Ecol Evol 20:23–27

Griffith SC, Owens IPF, Thuman KA (2002) Extra pair paternity in birds: a review of interspecific variation and adaptive function. Mol Ecol 11:2195–2212

Harrison AS (2014) The evolution and diversity of the Anolis dewlap. Dissertation, Harvard University

Hicks RM, Trivers RL (1983) The social behavior of Anolis valencienni. In: Rhodin GJ, Miyata K (eds) Recent Advances in Herpetology and Evolutionary Biology. Museum of Comparative Zoology, Cambridge, pp 570–595

Hinde RA (1956) The biological significance of the territories of birds. Ibis 98:340–369

Hrdy SB (1986) Empathy, polyandry, and the myth of the “coy” female. In: Bleier R (ed) Feminist Approaches to Science. Pergamon Press, New York, pp 119–146

Jenssen TA, Decourcy KR, Congdon JD (2005) Assessment in contests of male lizards (Anoliscarolinensis): how should smaller males respond when size matters? Anim Behav 69:1325–1336

Jenssen TA, Greenberg N, Hovde KA (1995) Behavioral profile of free-ranging male lizards, Anolis carolinensis, across breeding and post-breeding seasons. Herpetol Monogr 9:41–62

Jenssen TA, Nunez SC (1998) Spatial and breeding relationships of the lizard, Anolis carolinensis: Evidence of intrasexual selection. Behaviour 135:981–1003

Jenssen TA, Orrell KS, Lovern MB (2000) Sexual dimorphism in aggressive signal structure and use by a polygynous lizard, Anolis carolinensis. Copeia 2000:140–149

Johnson MA (2007) Behavioral ecology of Caribbean Anolis lizards: A comparative approach. Dissertation, Washington University

Kappeler PM, Barrett L, Blumstein DT, Clutton-Brock TH (2013) Constraints and flexibility in mammalian social behaviour: introduction and synthesis. Philos T Roy Soc B 368:0120337

Klug H (2011) Animal mating systems. eLS John Wiley and Sons Ltd Chichester

Losos JB (2009) Lizards in an Evolutionary Tree. University of California Press, Berkeley

Lott DF (1984) Intraspecific variation in the social systems of wild vertebrates. Behaviour 88:p266– 325

Maher CR, Lott DF (1995) Definitions of territoriality used in the study of variation in vertebrate spacing systems. Anim Behav 49:1581–1597

Martins EP (1994) Phylogenetic perspectives on the evolution of lizard territoriality. In: Vitt LJ, Pianka ER (eds) Lizard Ecology: Historical and Experimental Persepctives. Princeton University Press, Princeton, pp 117–144

Milam EL (2010) Looking for a few good males. The Johns Hopkins University Press, Baltimore

Noble GK, Bradley HT (1933) The mating behavior of lizards; its bearing on the theory of sexual selection. Ann NY Acad Sci 35:25–100

Oliver JA (1948) The anoline lizards of Bimini, Bahamas. Am Mus Novit 1383:1–36

Orians GH (1969) On the evolution of mating systems in birds and mammals. Am Nat 103:589–603

Orr TJ, Brennan PLR (2015) Sperm storage: distinguishing selective processes and evaluating criteria. Trends Ecol Evol 30:261–272

Passek KM (2002) Extra-pair paternity within the female-defense polygyny of the lizard, Anolis carolinensis: Evidence of alternative mating strategies. Dissertation, Virginia Polytechnic Institute

Philibosian R (1975) Territorial behavior and population regulation in the lizards, Anolis acutus and A. cristatellus. Copeia 1975:428–444

Qualls CP, Jaeger RG (1991) Dear enemy recognition in Anolis carolinensis. J Herpetol 25:361–363

Rand AS (1967a) Ecology and social organization in the iguanid lizard Anolis lineatopus. Proc US Nat Mus 122:1–79

Rand AS (1967b) The adaptive significance of territoriality in iguanid lizards. In: Milstead WW (ed) Lizard Ecology: A symposium. University of Missouri Press, Columbia, pp 106–115

Ruby DE (1984) Male breeding success and differential access to females in Anolis carolinensis. Herpetologica 40:272–280

Schoener TW (1981) An empirically based estimate of home range. Theor Popul Biol 20:281–325

Schoener TW, Schoener A (1980) Densities, sex ratios, and population structure in four species of Bahamian Anolis lizards. J Anim Ecol 49:19–53

Schoener TW, Schoener A (1982) Intraspecific variation in home-range size in some Anolis lizards. Ecology 63:809–823

Simon VB (2011) Communication signal rates predict interaction outcome in the brown anole lizard, Anolis sagrei. Copeia 2011:38–45

Stamps JA (1977). Social behavior and spacing patterns in lizards. In: Gans C, Tinkle DW (eds) Biology of the Reptilia, vol 7. Ecology and Behaviour A. Academic Press, New York, pp p265–334

Stamps JA (1983) Sexual selection, sexual dimorphism, and territoriality. In: Huey R, Pianka ER, Schoener TW (eds) Lizard Ecology. Harvard University Press, Cambridge, pp 169–204

Stamps JA (1994) Territorial behavior: Testing the assumptions. Adv Stud Behav 23:173–231

Stamps JA (1995) Using growth-based models to study behavioral factors affecting sexual size dimorphism. Herpetol Monogr 9:75–87

Tang-Martinez Z (2016) Rethinking Bateman’s principles: Challenging persistent myths of sexually reluctant females and promiscuous males. J Sex Res 53:532–559

Tang-Martinez Z, Ryder TB (2005) The problem with paradigm: Bateman’s worldview as a case study. Integr Comp Biol 45:821–830

Tewksbury JJ, Anderson JGT, Bakker JD et al (2014) Natural history’s place in science and society. Bioscience 64:300–310

Thompson FG (1954) Notes on the behavior of the lizard Anolis carolinensis. Copeia 1954:299

Tinbergen N (1957) The functions of territory. Bird Study 4:14–27

Tokarz RR (1998) Mating pattern in the lizard Anolis sagrei: Implications for mate choice and sperm competition. Herpetologica 54:388–394

Tokarz RR (2008) Males distinguish between former female residents of their territories and unfamiliar nonresident females as preferred mating partners in the lizard Anolis sagrei. J Herpetol 42:260–264

Tokarz RR, McMann S, Seitz L, John-Alder H (1998) Plasma corticosterone and testosterone levels during the annual reproductive cycle of male brown anoles (Anolis sagrei). Physiol Zool 71:139–146

Trivers RL (1976) Sexual selection and resource-accruing abilities in Anolis garmani. Evolution 30:253–269

Uller T, Olsson M (2008) Multiple paternity in reptiles: patterns and processes. Mol Ecol 17:p2566–2580

Webster R, Greenberg N (1988) Territoriality and social dominance in the green anole lizard. Am Zool 28:A73

## References

1. Alworth, T.J. 1986. Perch availability and season affect aggression levels in the territorial lizard, Anolis carolinensis. American Zoologist 26: 1041.

2. Andrews, R.M. 1985. Male choice by females of the lizard, Anolis carolinensis. Journal of Herpetology 19: 284–289.

3. Brach, V. 1976. Habits and food of Anolis equestris in Florida. Copeia 1976: 187–189.

4. Bull, C.M. 2000. Monogamy in lizards. Behavioural Processes 51: 7–20.

5. Bush, J.M., M.M. Quinn, E.C. Balreira, M.A. Johnson. 2016. How do lizards determine dominance? Applying ranking algorithms to animal social behaviour. Animal Behaviour 118: 65–74.

6. Calsbeek, R. 2009. Sex-specific adult dispersal and its selective consequences in the brown anole, Anolis sagrei. Journal of Animal Ecology 78: 617–624.

7. Calsbeek, R., C. Bonneaud, S. Prabhu, N. Manoukis, and T.B. Smith. 2007. Multiple paternity and sperm storage lead to increased genetic diversity in Anolis lizards. Evolutionary Ecology Research 9: 495–503.

8. Calsbeek, R., W. Buermann, and T.B. Smith. 2009. Parallel shifts in ecology and natural selection in an island lizard. BMC Evolutionary Biology 9: 3.

9. Calsbeek, R., and E. Marnocha. 2006. Context dependent territory defense: the importance of habitat structure in Anolis sagrei. Ethology 112: 537–543.

10. Calsbeek, R., and T.B. Smith. 2007. Probing the adaptive landscape using experimental islands: density-dependent natural selection on lizard body size. Evolution 61: 1052–1061.

11. Carpenter, C.R. 1958. Territoriality: A review of concepts and problems. Pp. 224–250 In A. Roe, and G.G. Simpson (eds.) Behavior and Evolution. Yale University Press, New Haven, CT, USA.

12. Charles, G.K., and T.J. Ord. 2012. Factors leading to the evolution and maintenance of a male ornament in territorial species. Behavioral Ecology and Sociobiology 66: 231–239.

13. Colnaghi, G.L. 1971. Partitioning of a restricted food source in a territorial iguanid (Anolis carolinensis). Psychonomic Science 23: 59–60.

14. Crews, D. 1980. Studies in squamate sexuality. Bioscience 30: 835–838.

15. Crews, D., and N. Greenberg. 1981. Function and causation of social signals in lizards. American Zoologist 21: 273–294.

16. Decourcy, K.R., and T.A. Jenssen. 1994. Structure and use of male territorial headbob signals by the lizard Anolis carolinensis. Animal Behaviour 47: 251–262.

17. Eason, P.K., and J.A. Stamps. 1992. The effect of visibility on territory size and shape. Behavioral Ecology 3: 166–172.

18. Evans, L.T. 1936a. Social behavior of the normal and castrated lizard, Anolis carolinensis. Science 83: 104.

19. Evans, L.T. 1936b. Territorial behavior of normal and castrated females of Anolis carolinensis. Pedagogical Seminary and Journal of Genetic Psychology 49: 49–60.

20. Evans, L.T. 1936c. A study of a social hierarchy in the lizard Anolis carolinensis. Pedagogical Seminary and Journal of Genetic Psychology 48: 88–111.

21. Evans, L.T. 1938a. Cuban field studies on the territoriality of the lizard Anolis sagrei. Journal of Comparative Psychology 25: 97–125.

22. Evans, L.T. 1938b. Courtship behavior and sexual selection of Anolis. Journal of Comparative Zoology 26: 475–497.

23. Everly, A., L.M. Sievert, and R.B. Thomas. 2011. Dear enemy recognition in captive brown anoles. Journal of Kansas Herpetology 40: 13–16.

24. Farrell, W.J., and W. Wilczynski. 2006. Aggressive experience alters place preference in green anole lizards, Anolis carolinensis. Animal Behaviour 71: 1155–1164.

25. Fitch, H.S. 1976. Sexual size differences in the mainland anoles. Occasional papers of the Museum of Natural History, the University of Kansas 50: 1–21.

26. Fitch, H.S., and R.W. Henderson. 1976. A field study of the rock anoles (Reptilia, Lacertilia, Iguanidae) of Southern Mexico. Journal of Herpetology 10: 303–311.

27. Fitch, H.S., and R.W. Henderson. 1987. Ecological and ethological parameters in Anolis bahorucoensis, a species having rudimentary development of the dewlap. Amphibia-Reptilia 8: 69–80.

28. Flanagan, S.P., and C.R. Bevier. 2014. Do male activity level and territory quality affect female association time in the brown anole, Anolis sagrei? Ethology 120: 365–374.

29. Fleishman, L.J. 1988. The social behavior of Anolis auratus, a grass anole from Panama. Journal of Herpetology 22: 13–23.

30. Fobes, T.M., R. Powell, J.S. Parmerlee Jr., A. Lathrop, and D.D. Smith. 1992. Natural history of Anolis cybotes (Sauria: Polychridae) from an altered habitat in Barahona, Dominican Republic. Caribbean Journal of Science 28: 200–207.

31. Forster, G.L., M.J. Watt, W.J. Korzan, K.J. Renner, and C.H. Summers. 2005. Opponent recognition in male green anoles, Anolis carolinensis. Animal Behaviour 69: 733–740.

32. Gordon, R.E. 1956. The biology and biodemography of Anolis carolinensis carolinensis Voight. Ph.D. Dissertation, Tulane University.

33. Gorman, G.C. 1969. Intermediate territorial display of a hybrid Anolis lizard (Sauria: Iguanidae). Zeitschrift für Tierpsychologie 26: 390–393.

34. Greenberg, B., and G.K. Noble. 1944. Social behavior of the American chameleon (Anolis carolinensis Voight). Physiological Zoology 17: p392–439.

35. Henningsen, J.P., and D.J. Irschick. 2012. An experimental test of the effect of signal size and performance capacity on dominance in the green anole lizard. Functional Ecology 26: 3–10.

36. Harrison, A.S. 2014. The evolution and diversity of the Anolis dewlap. Ph.D. Dissertation, Harvard University.

37. Hicks, R.M., and R.L. Trivers. 1983. The social behavior of Anolis valencienni. Pp. 570–595 In G.J. Rhodin, and K. Miyata. Recent Advances in Herpetology and Evolutionary Biology. Museum of Comparative Zoology, Cambridge, MA, USA.

38. Jackson, J.F. 1973. Notes on the population biology of Anolis tropidonotus in a Honduran highland pine forest. Journal of Herpetology 7: 309–311.

39. Jenssen, T.A. 1970. The ethoecology of Anolis nebulosus (Sauria, Iguanidae). Journal of Herpetology, 4: 1–38.

40. Jenssen, T.A. 2002. Spatial awareness by the lizard Anolis cristatellus: why should a non-ranging species demonstrate homing behavior? Herpetologica 58: 364–371.

41. Jenssen, T.A., K.R. Decourcy, and J.D. Congdon. 2005. Assessment in contests of male lizards (Anolis carolinensis): how should smaller males respond when size matters? Animal Behaviour 69: 1325–1336.

42. Jenssen, T.A., and P.C. Feeley. 1991. Social behavior of the male anoline lizard Chamaelinorops barbouri, with a comparison to Anolis. Journal of Herpetology 25: 454–462.

43. Jenssen, T.A, N. Greenberg, and K.A. Hovde. 1995. Behavioral profile of free-ranging male lizards, Anolis carolinensis, across breeding and post-breeding seasons. Herpetological Monographs 9: 41–62.

44. Jenssen, T.A., M.B. Lovern, J.D. Congdon. 2001. Field-testing the protandry based mating system for the lizard, Anolis carolinensis: does the model organism have the right model? Behavioral Ecology and Sociobiology 50: 162–172.

45. Jenssen, T.A., and S.C. Nunez. 1994. Male and female reproductive cycles of the Jamaican lizard, Anolis opalinus. Copeia 1994: 767–780

46. Jenssen, T.A., and S.C. Nunez. 1998. Spatial and breeding relationships of the lizard, Anolis carolinensis: Evidence of intrasexual selection. Behaviour 135: 981–1003.

47. Jenssen, T.A., K.S. Orrell, and M.B. Lovern. 2000. Sexual dimorphism in aggressive signal structure and use by a polygynous lizard, Anolis carolinensis. Copeia 2000: 140–149.

48. Jiménez, R.R., and J.A. Rodríguez-Rodríguez. 2015. The relationship between perch type and aggressive behavior in the lizard Norops polylepis (Squamata: Dactyloidae). Phyllomedusa 14: 43–51.

49. Johnson, M.A., L.J. Revell, and J.B. Losos. 2010. Behavioral convergence and adaptive radiation: effects of habitat use on territorial behavior in Anolis lizards. Evolution 64: p1151–1159.

50. Johnson, M.A. 2007. Behavioral ecology of Caribbean Anolis lizards: A comparative approach. Ph.D. Dissertation, Washington University.

51. Joyce, T., D.A. Eifler, and R. Powell. 2010. Variable habitat use influences the mating system of a Lesser Antillean anole. Amphibia-Reptilia 31: 395–401.

52. Kaiser, B.W., and H.R. Mushinsky. 1994. Tail loss and dominance in captive adult male Anolis sagrei. Journal of Herpetology 28: 342–346.

53. Lailvaux, S.P., and D.J. Irschick. 2007. The evolution of performance-based male fighting ability in Caribbean Anolis lizards. The American Naturalist 170: 573–586.

54. Leuck, B.E. 1995. Territorial defense by male green anoles: An experimental test of the roles of residency and resource quality. Herpetological Monographs 9: 63–74.

55. Lister, B.C. 1976. The nature of niche expansion in West Indian Anolis lizards II: Evolutionary components. Evolution 30: 677–692.

56. Lovern, M.B. 2000. Behavioral ontogeny in the free-ranging juvenile male and female green anoles, Anolis carolinensis, in relation to sexual selection. Journal of Herpetology 34: 274–281.

57. McMann, S. 2000. Effects of residence time on displays during territory establishment in a lizard. Animal Behaviour 59: 513–522.

58. McMann, S., and A.V. Paterson. 2003. The relationship between location and displays in a territorial lizard. Journal of Herpetology 37: 414–416.

59. McMann, S., and A.V. Paterson. 2012. Display behavior of resident brown anoles (Anolis sagrei) during close encounters with neighbors and non-neighbors. Herpetological Conservation and Biology 7: 27–37.

60. Nicholson, K.E., and P.M. Richards. 2011. Home-range size and overlap within an introduced population of the Cuban knight anole, Anolis equestris (Squamata: Iguanidae). Phyllomedusa 10:65–73.

61. Noble, G.K., and H.T. Bradley. 1933. The mating behavior of lizards; its bearing on the theory of sexual selection. Annals of the New York Academy of Science 35: 25–100.

62. Nunez, S.C., T.A. Jenssen, and K. Ersland. 1997. Female activity profile of a polygynous lizard (Anolis carolinensis): Evidence of intersexual asymmetry. Behaviour 134: 205–223.

63. Oliver, J.A. 1948. The anoline lizards of Bimini, Bahamas. American Museum Novitates 1383: 1–36.

64. Orrell, K.S., and T.A. Jenssen. 2002. Male mate choice by the lizard Anolis carolinensis: a preference for novel females. Animal Behaviour 63: 1091–1102.

65. Orrell, K.S., and T.A. Jenssen. 2003. Heterosexual signalling by the lizard Anolis carolinensis, with intersexual comparisons across contexts. Behaviour 140: 603–634.

66. Passek, K.M. 2002. Extra-pair paternity within the female-defense polygyny of the lizard, Anolis carolinensis: Evidence of alternative mating strategies. Ph.D. Dissertation, Virginia Polytechnic Institute.

67. Paterson, A.V. 1999. Effects of prey availability on perch height of female bark anoles, Anolis distichus. Herpetologica 55: 242–247.

68. Paterson, A.V. 2002. Effects of an individual’s removal on space use and behavior in territorial neighborhoods of brown anoles (Anolis sagrei). Herpetologica 58: 382–393.

69. Paterson, A.V., and S. McMann. 2004. Differential headbob displays toward neighbors and nonneighbors in the territorial lizard Anolis sagrei. Journal of Herpetology 38: 288–291.

70. Pereira, H.M., S.R. Loarie, and J. Roughgarden. 2002. Monogamy, polygyny and interspecific interactions in the lizards Anolis pogus and Anolis gingivinus. Caribbean Journal of Science 38: 132–136.

71. Philibosian, R. 1975. Territorial behavior and population regulation in the lizards, Anolis acutus and A. cristatellus. Copeia 1975: 428–444.

72. Qualls, C.P. and R.G. Jaeger. 1991. Dear enemy recognition in Anolis carolinensis. Journal of Herpetology 25: 361–363.

73. Rand, A.S. 1967a. Ecology and social organization in the iguanid lizard Anolis lineatopus. Proceedings of the United States National Museum 122: 1–79.

74. Rand, A.S. 1967b. The adaptive significance of territoriality in iguanid lizards. Pp. 106–115 In Lizard Ecology: A symposium. University of Missouri Press, Columbia, MO, USA.

75. Reagan, D.P. 1992. Congeneric species distribution and abundance in a three-dimensional habitat: the rain forest anoles of Puerto Rico. Copeia 1992: 392–403.

76. Ruby, D.E. 1984. Male breeding success and differential access to females in Anolis carolinensis. Herpetologica 40: 272–280.

77. Ruibal, R., and R. Philibosian. 1974a. The population ecology of the lizard Anolis acutus. Ecology 55: 525–537.

78. Ruibal, R., and R. Philibosian. 1974b. Aggression in the lizard Anolis acutus. Copeia 1974: p349–357.

79. Schoener, T.W. 1981. An empirically based estimate of home range. Theoretical Population Biology 20: 281–325.

80. Schoener, T.W., and A. Schoener. 1980. Densities, sex ratios, and population structure in four species of Bahamian Anolis lizards. Journal of Animal Ecology 49: 19–53.

81. Schoener, T.W., and A. Schoener. 1982. Intraspecific variation in home-range size in some Anolis lizards. Ecology 63: 809–823.

82. Schoener, T.W., and A. Schoener. 1982. The ecological correlates of survival in some Bahamian Anolis lizards. Oikos 39: 1–16.

83. Sexton, O.J., H.F. Heatwole, and E.H. Meseth. 1963. Seasonal population changes in the lizard, Anolis limifrons, in Panama. The American Midland Naturalist 69: 482–491.

84. Simon, V.B. 2011. Communication signal rates predict interaction outcome in the brown anole lizard, Anolis sagrei. Copeia 2011: 38–45.

85. Stamps, J.A. 1973. Displays and social organization in female Anolis aeneus. Copeia 1973: p264–272.

86. Stamps, J.A. 1977a. The relationship between resource competition, risk, and aggression in a tropical territorial lizard. Ecology 58: 349–358.

87. Stamps, J.A. 1977b. Social behavior and spacing patterns in lizards. Pp. 265–334 In C. Gans, and D.W. Tinkle (eds.) Biology of the Reptilia. Volume 7. Ecology and Behaviour A. Academic Press Inc., New York, NY, USA.

88. Stamps, J.A. 1978. A field study of the ontogeny of social behavior in the lizard Anolis aeneus. Behaviour 66: 1–31.

89. Stamps, J.A. 1983. Sexual selection, sexual dimorphism, and territoriality. Pp. 169–204 In R. Huey, E.R. Pianka, and T.W. Schoener (eds.) Lizard Ecology. Harvard University Press, Cambridge, MA, USA.

90. Stamps, J.A. 1994. Territorial behavior: Testing the assumptions. Advances in the Study of Behavior 23: 173–231.

91. Stamps, J.A. 1995. Using growth-based models to study behavioral factors affecting sexual size dimorphism. Herpetological Monographs 9: 75–87.

92. Stamps, J.A. 1999. Relationships between female density and sexual size dimorphism in samples of Anolis sagrei. Copeia 1999: 760–765.

93. Stamps, J.A. 2001. Learning from lizards. Pp. 149–168 In L.A. Dugatkin (ed.) Model Systems in Behavioral Ecology: Integrating Conceptual, Theoretical and Empirical Approaches. Princeton University Press, Princeton, NJ, USA.

94. Stamps, J.A., J.B. Losos, and R.M. Andrews. 1997. A comparative study of population density 977 and sexual size dimorphism in lizards. The American Naturalist 149: 64–90.

95. Stamps, J.A., V.V. Krishnan, and R.M. Andrews. 1994. Analysis of sexual size dimorphism using null growth-based models. Copeia 1994: 598–613.

96. Talbot, J.J. 1979. Time budget, niche overlap, inter- and intraspecific aggression in Anolis humilis and A. limifrons from Costa Rica. Copeia 1979: 472–481.

97. Thompson, F.G. 1954. Notes on the behavior of the lizard Anolis carolinensis. Copeia 1954: 299.

98. Tokarz, R.R. 1985. Body size as a factor determining dominance in staged agonistic encounters between male brown anoles (Anolis sagrei). Animal Behaviour 33: 746–753.

99. Tokarz, R.R. 1995. Mate choice in lizards: A review. Herpetological Monographs 9: 17–40.

100. Tokarz, R.R. 1998. Mating pattern in the lizard Anolis sagrei: Implications for mate choice and sperm competition. Herpetologica 54: 388–394.

101. Tokarz, R.R. 2008. Males distinguish between former female residents of their territories and unfamiliar nonresident females as preferred mating partners in the lizard Anolis sagrei. Journal of Herpetology 42: 260–264.

102. Tokarz, R.R., S. McMann, L. Seitz, and H. John-Alder. 1998. Plasma corticosterone and testosterone levels during the annual reproductive cycle of male brown anoles (Anolis sagrei). Physiological Zoology 71: 139–146.

103. Tokarz, R.R., A.V. Paterson, and S. McMann. 2003. Laboratory and field test of the functional significance of the male’s dewlap in the lizard Anolis sagrei. Copeia 2003: 502–511.

104. Trivers, R.L. 1976. Sexual selection and resource-accruing abilities in Anolis garmani. Evolution 30: 253–269.

105. Webster, R. and N. Greenberg. 1988. Territoriality and social dominance in the green anole lizard. American Zoologist 28: A73.

106. Yang, E., S.M. Phelps, D. Crews, and W. Wilcynski. 2001. The effects of social 1001 experience on aggressive behavior in the green anole lizard (Anolis carolinensis). Ethology 107: 777–793.

